# Arrest of mouse preterm labor until term delivery by combination therapy with atosiban and mundulone, a natural product with tocolytic efficacy

**DOI:** 10.1101/2023.06.06.543921

**Authors:** Shajila Siricilla, Christopher J. Hansen, Jackson H. Rogers, Debasmita De, Carolyn L. Simpson, Alex G. Waterson, Gary A. Sulikowski, Stacey L. Crockett, Naoko Boatwright, Jeff Reese, Bibhash C. Paria, J Newton, Jennifer L. Herington

**Affiliations:** Division of Neonatology, Department of Pediatrics, Vanderbilt University Medical Center, Nashville, TN, USA; Department of Pharmacology, Vanderbilt University, Nashville, TN, USA; Vanderbilt Institute of Chemical Biology, Vanderbilt University, Nashville, TN, USA; Division of Maternal Fetal Medicine, Department of Obstetrics and Gynecology, Vanderbilt University Medical Center, Nashville, TN, USA

**Keywords:** drug combination, mundulone, natural products, preterm labor, preterm birth, tocolytic, uterus

## Abstract

Currently, there is a lack of FDA-approved tocolytics for the management of preterm labor (PL). In prior drug discovery efforts, we identified mundulone and its analog mundulone acetate (MA) as inhibitors of *in vitro* intracellular Ca^2+^-regulated myometrial contractility. In this study, we probed the tocolytic and therapeutic potential of these small molecules using myometrial cells and tissues obtained from patients receiving cesarean deliveries, as well as a mouse model of PL resulting in preterm birth. In a phenotypic assay, mundulone displayed greater efficacy in the inhibition of intracellular-Ca^2+^ from myometrial cells; however, MA showed greater potency and uterine-selectivity, based IC_50_ and E_max_ values between myometrial cells compared to aorta vascular smooth muscle cells, a major maternal off-target site of current tocolytics. Cell viability assays revealed that MA was significantly less cytotoxic. Organ bath and vessel myography studies showed that only mundulone exerted concentration-dependent inhibition of *ex vivo* myometrial contractions and that neither mundulone or MA affected vasoreactivity of ductus arteriosus, a major fetal off-target of current tocolytics. A high-throughput combination screen of *in vitro* intracellular Ca^2+^-mobilization identified that mundulone exhibits synergism with two clinical-tocolytics (atosiban and nifedipine), and MA displayed synergistic efficacy with nifedipine. Of these synergistic combinations, mundulone + atosiban demonstrated a favorable *in vitro* therapeutic index (TI)=10, a substantial improvement compared to TI=0.8 for mundulone alone. The *ex vivo* and *in vivo* synergism of mundulone and atosiban was substantiated, yielding greater tocolytic efficacy and potency on isolated mouse and human myometrial tissue and reduced preterm birth rates in a mouse model of PL compared to each single agent. Treatment with mundulone 5hrs after mifepristone administration (and PL induction) dose-dependently delayed the timing of delivery. Importantly, mundulone in combination with atosiban (FR 3.7:1, 6.5mg/kg + 1.75mg/kg) permitted long-term management of PL after induction with 30 μg mifepristone, allowing 71% dams to deliver viable pups at term (> day 19, 4-5 days post-mifepristone exposure) without any visible maternal and fetal consequences. Collectively, these studies provide a strong foundation for the future development of mundulone as a stand-alone single- and/or combination-tocolytic therapy for management of PL.

## 1. Introduction

Preterm birth (PTB) rates continue to rise with over 15 million cases/year globally and remains the greatest contributor to neonatal morbidities and mortalities [1, 2]. The causes of PTB are multifactorial, yet in most situations, PTB follows spontaneous preterm labor (sPL) without a known maternal, fetal, or placental condition. Regardless of the trigger, the common denominator for all known causes of PTB is the early activation of uterine contractions, which occurs through stimulation of intracellular Ca^2+^-release in myometrial cells [3, 4]. The American College of Obstetricians and Gynecologists’ Committee on Practice Bulletins [5], World Health Organization [6] and nine other clinical practice guidelines [7] agree on the short term tocolytic benefit for PTB. Moreover, outcome studies consistently show the substantial neonatal benefits for each additional week *in utero* [8–12].

Spontaneous PL is typically managed with off-label use of tocolytic drugs including: nifedipine (calcium channel blocker), indomethacin (cyclooxygenase inhibitor) or terbutaline (beta-adrenergic agonist). However, these drugs are not FDA-approved for tocolytic use due to detrimental off-target side effects and/or short duration of benefit. Specifically, these drugs are known to cause either maternal cardiovascular effects, *in utero* constriction of the fetal ductus arteriosus (DA) or fetal tachycardia [13–16]. These side effects are not surprising, given that these drugs were initially developed to treat disorders in other smooth muscle types such as vascular and pulmonary. Drug development for uterospecific drugs has yielded (i) oxytocin-receptor (OXTR) antagonists, such as atosiban, for tocolytic use and (ii) a synthetic progestin (17-alpha hydroxyprogesterone caproate, ‘Makena™’) for the prevention of sPL. However, FDA-approval for Atosiban has been denied due to controversy concerning its effectiveness in clinical trials to delay PTB and improve neonatal outcomes [17–19]. Furthermore, Makena’s manufacturer recently volunteered to withdraw the drug from the US market in response to FDA recommendations based on recent large-scale studies showing less significant reductions in sPL [20]. Thus, novel safe and effective tocolytics are urgently needed for management of sPL.

Combination therapy offers an opportunity to increase the efficacy or potency due to the additive or synergistic effect, thereby decreasing the toxicity or side effects associated with higher doses of single drugs. Past studies examining the benefit of combined tocolytic therapy only examined a single dose of each drug, without determining the optimal fixed ratio required to achieve synergy [21–26]. Thus, while combinations of current tocolytics ranked highest for delaying preterm birth by 48 hours and 7 days, they also result in more frequent cessation due to adverse effects [27].

In our efforts to discover novel tocolytics, we previously performed a high-throughput screen against a library of small molecules comprised of drug components, natural products, and other bioactive molecules with a wide range of biological activity [3]. We identified the isoflavone natural-product mundulone, and its derivative mundulone acetate (MA), as potent antagonists of Ca^2+^-mobilization from primary myometrial cells.

Mundulone is extracted from the bark of *Mundulea sericea* and its structure was elucidated in 1959 as part of the isoflavone subclass of flavonoids [28]. Isoflavones have several pharmacological effects, including modulation of smooth muscle contractions in the uterus and vasculature [29–33]. Moreover, a recent review on natural products for tocolysis found that most plant extracts for inhibition of uterine contractions belong to flavonoid or terpene classes [34], with *Bryophyllum pinnatum* entering into a clinical trial for preterm labor, which was withdrawn early due to lack of patient enrollment [35–40]. Therefore, mundulone and MA belong to a promising class of compounds to study for tocolytic potential.

Based on our prior discovery that mundulone and MA inhibit *in vitro* myometrial Ca^2+^-mobilization and the reported tocolytic ability of other isoflavones, we aimed to further probe mundulone and MA for tocolytic drug development. The goal of this study was to: 1) examine the uterineLselectivity of Ca^2+^-mobilization inhibition, 2) determine cytotoxicity, 3) identify synergistic combinations of mundulone and MA with current clinically-utilized tocolytics, 4) establish the *ex vivo* tocolytic efficacy and potency and confirm uterine selectivity at the tissue level and 5) evaluate the *in vivo* efficacy to delay delivery onset using a mouse model of PL and PTB.

## 2. Materials and methods

### 2.1. Primary human uterine myometrial tissue and cells

Human myometrial biopsies were obtained at the time of cesarean section under the Vanderbilt University Medical Center Institutional Review Board protocol #150791 and according to The Code of Ethics of the World Medical Association (Declaration of Helsinki). Reproductive-age (18-45 years) women undergoing a scheduled or repeat cesarean section at term gestation (≥39 weeks) were recruited, fully informed, and consented to the study. Inclusion criteria included breech presentation, previous cesarean section, fetal anomaly, fetal distress or placenta previa. Exclusion criteria included: clinical or histological signs of vaginal/chorioamniotic/intrauterine inflammation/infection, active SARS-CoV-2, HIV, Hepatitis B, current use of vasopressors or bronchodilators and insulin-controlled diabetes mellitus. Samples were placed into sterile cold 1XHBSS or 1XPBS and transported to the laboratory for immediate myometrial cell isolation or use in *ex vivo* organ bath studies, respectively.

### 2.2 Cell culture

Primary myometrial cells were isolated, selectively enriched for smooth muscle cells and cultured in complete media (DMEM supplemented with 10% FBS, 25 mM HEPES, 100 U/ml penicillin-streptomycin) as previously described [3, 41]. Unpassaged cells became near-confluent 10-15 days post-isolation. Human primary aortic vascular smooth muscle cells (VSMCs; ATCC PCS-100-012) were cultured in complete vascular cell growth medium and growth kit (ATCC PCS-100-030 and ATCC PCS-100-042). Myometrial and VSMCs were plated at 4,000 cells/well in black-walled 384-well plates (Grenier Bio-One) for Ca^2+^-mobilization assays exactly as previously described [41].

### 2.3. Comparative concentration-dependent Ca^2+^-mobilization

Mundulone and MA (MicroSource Discovery Systems 00200011 and 00200019) were tested at 10-point three-fold titrations starting at 60 μM in triplicate to compare the E_max_ and IC_50_ values between myometrial and VSMCs in a Ca^2+^-mobilization assay, performed as previously described. [3, 41]. Data were analyzed using WaveGuide software and the % inhibition for each concentration of mundulone and MA was calculated.

### 2.4. Combination high-throughput Ca^2+^-mobilization assay

An 8X8 dose matrix was used to evaluate the combination effects between two serially diluted single-compound concentrations. Mundulone and MA were combined with atosiban (Sigma A3480), indomethacin (Cayman Chemicals 70270), or nifedipine (Cayman Chemicals 11106). The single-compound IC_50_ values were utilized with three 2-fold titrations above and below the IC_50_ value. Control (no compound) additions were included in the dose matrix. Moreover, the first row and the first column of each matrix contained the individual compound concentrations used to compare the effect of combination. The high-throughput Ca^2+^-assay was performed in 384-well format, allowing for 6 combinations per plate, using the methods described above.

Precise titrations of each compound were performed using an Echo555 instrument within the Vanderbilt High-Throughput Screening Facility. Raw data was first analyzed using Waveguide, as described above and previously reported [41]. After calculating the % response for each compound concentration examined, synergistic combinations were determined using Combenefit software [42], which provided synergy scores and heat-maps to visualize models of Bliss independence, Highest Single Agent, and Loewe additivity. After selecting up to 3 fixed ratios (FRs) of synergistic drug combinations, a concentration-response analysis was performed as described in detail above, to confirm whether the synergy was a result of increased efficacy and/or potency.

### 2.5. Cell viability assay

Cytotoxicity of single- and combination-compounds on myometrial (hTERT-immortalized human myometrial, hTERT-HM) [41, 43, 44], kidney (human primary renal proximal tubule epithelial cells, RPTEC), and liver (human hepatocellular carcinoma, HepG2) [45–50] cells were assessed using a standard WST-1 (Roche) cell viability assay. hTERT-HM (kindly provided by Dr. Jennifer Condon, Wayne State University), RPTEC (ATCC PCS-400-010), and HepG2 (ATCC HB-8065) cells were cultured in complete DMEM/F-12 (Gibco 11- 039-02), RPTEC media (ATCC PCS-400-030) and EMEM (ATCC 30-2003), respectively. DMEM/F-12 and HEPG2 media were supplemented with 10% FBS and 1% penicillin-streptomycin, while RPTEC media was supplemented with growth kit components (ATCC PCS-400-040). Two-fold serial dilutions of each single-compound were tested at 0.78–200 μM (a final volume of 100μL). Additionally, compound combinations at their synergistic FR and matched single-compound concentrations were tested, as were negative (DMSO) and positive (napabucasin) controls. Cells (4 x 10^4^ per well) in 96-well culture plates were incubated with compounds for 72hrs, after which 10 μL of WST-1 was added to each well. Absorbance at 450nm and 600nm was read 2hrs after incubation with WST-1. After deduction of background absorbance, the % inhibition in cell viability was calculated. An *in vitro* therapeutic index (TI) was calculated as a ratio of the smallest IC_50_ value among the three cell types (concentration of the compound required to affect 50% of cell viability) to the IC_50_ value from the Ca^2+^ mobilization assay [51, 52].

### 2.6. Mouse tissue sample collection

Animal experiments were approved by the Vanderbilt University Institutional Animal Care and Use Committee and conformed to the guidelines established by National Institutes of Health for the care and use of Laboratory animals. Adult (8–12wk) female CD-1 IGS (Charles River Laboratories) mice were bred with fertile males in a restricted 3hr window (8-11am), and the presence of a copulatory plug was considered day 1 of pregnancy. The average date of delivery for these mice in our colony occurs on day 19.5. Mice were euthanized with an overdose of isoflurane, followed by cervical dislocation. The uterus was excised on the morning of day 19 of pregnancy to obtain myometrial strips for *ex vivo* contractility studies, as well as the collection of fetal DA tissue for *ex vivo* myography experiments described below.

### 2.7. *Ex vivo* myometrial contractility assay

An isometric contractility organ bath assay was performed as previously described [3, 53]. Briefly, mouse and human myometrial strips (12mm X 5mm X 1 mm, N= 6-11 from 4-5 mice and 8 humans) were submerged in a heated and oxygenated (37°C, 95% O2-5% CO2) Radnoti LLC tissue bath containing Kreb’s Bicarbonate Solution. Each mouse strip was placed under 1g tension and allowed to equilibrate in the organ bath for 60min prior to recording baseline spontaneous contractile activity. Human tissue was placed under 3g of tension, washed with Kreb’s Bicarbonate Solution and treated with KCl, as previously described [54]. Rhythmic contractions were typically established within ∼2-3 hours.

Stock 0.1M mundulone and MA were dissolved in 100% PEG-400, while 0.027M of atosiban was also dissolved in PEG-400. Following the establishment of rhythmic, spontaneous contractions, cumulative concentrations (1nM to 0.1mM) of mundulone, MA, or vehicle control (PEG-400) were added to individual organ baths every 15 or 20 min depending on whether mouse or human tissue was utilized for the experiment, respectively. In a second set of experiments, cumulative concentrations (1nM to 0.1mM) of a mundulone + atosiban combination at a FR 3.7:1, as well as mundulone and atosiban alone (at their respective concentrations reflective of the ratio) were added to individual organ baths. After the highest concentration of drug or vehicle examined, tissue viability at the end of the experiment was evaluated using exposure to 75mM KCl.

Isometric contractions were recorded using PowerLab/8 SP (ADInstruments) equipment and analyzed with LabChart 7 Pro software (ADInstruments). Contractile activity was assessed by AUC/duration (integral relative to baseline divided by the duration of each treatment period assessed). All treatment data were expressed as a percent change of baseline spontaneous contractile activity.

### 2.8. *Ex vivo* fetal DA myography

The vasoreactivity of fetal mouse DAs was evaluated using cannulated, pressurized vessel myography and computer-assisted videomicroscopy, as previously described [55, 56]. Briefly, the ductus vessels (7–9 fetal mice, representing at least three different litters) were mounted in custom myography chambers placed on inverted microscopes equipped with a digital image capture system (IonOptix) to record changes in the intraluminal vessel diameter. The pressure was increased in 5-mmHg increments to 20mmHg, followed by treatment with 50mM KCl deoxygenated modified Krebs buffer to determine vessel viability and peak contractility. The vessels were then changed from a flow-through system to a 20ml recirculating system and allowed to equilibrate for an additional 20 minutes. Baseline lumen diameter was recorded prior to adding cumulative concentrations (1nM to 0.1mM) of mundulone, MA, or vehicle control every 20 mins. After the highest concentration of drug or vehicle examined, vessel viability was determined via exposure to 50mM KCl. All treatment data were expressed as a percent change in baseline lumen diameter.

### 2.9. *In vitro* ADME assays

Mundulone and MA were submitted to Q^2^ Solutions for Tier 1 screening assays (metabolic stability, permeability and CYP inhibition) according to standard protocols.

### 2.10. *In vivo* therapeutic efficacy in mifepristone-induced mouse model of PTB

CD-1 female and male mice were bred in a restricted 3hr window (8-11am), and the presence of a copulatory plug was considered day 1 of pregnancy. On day 15 of pregnancy, mice (N≥5 per group) were administered (s.c.) various doses of mifepristone (10, 20, 25, 30, 40, 60, 80, 100, 150 μg; Sigma M8046) to establish the lowest dose required to induce PTB in 100% mice (prior to day 18.5 of pregnancy, which is 24hrs prior to normal term delivery). Based on the results of these studies, day 15 pregnant mice (average weight: 44.97 ± 4.35g) were administered either 30 μg or 150 μg (commonly-utilized dose) mifepristone, followed by administration (s.c.) of treatment 5hrs after the onset of *in vivo* uterine contractions [57]. Treatments included either vehicle (0.7mL/kg ethanol in 150μL sesame oil, negative control), 5mg/kg nifedipine (positive control), mundulone alone (6.5, 13, or 26 mg/kg), atosiban alone (1.76 or 3.5 mg/kg), or mundulone + atosiban (6.5+1.76 or 13+3.5 mg/kg). Mice (N≥5 per group) received twice daily (8am and 8pm) treatments of vehicle or test molecules on days 16-18. The onset of delivery was monitored using infrared video surveillance and defined as delivery of the first pup. Preterm birth was defined as delivery occurring 24 hrs prior to the average onset of term delivery of untreated mice. The number of pups and neonatal outcome (surviving pups born at term per total litter size including non-viable pups born, pups resorbed *in utero* and pup weights at term delivery) were recorded.

### 2.11. Statistical analysis

Data were analyzed using GraphPad Prism software and are expressed as mean ± SEM. Non-linear regression analyses were performed to generate CRCs for calculation of IC_50_ and E_max_. Comparisons of fit determined whether three-parameter non-linear log fit lines were significantly different between: 1) mundulone, MA, and the vehicle control, as well as 2) mundulone alone, atosiban alone, and the mundulone + atosiban combination. Two-way analysis of variance followed by a *post hoc* Tukey test for multiple comparisons was used to determine significant differences between the % response for each concentration of compound. Two-way analysis of variance followed by a post hoc Fisher’s LSD test was used to determine significant differences between the E_max_ values. Ordinary one-way ANOVA was performed to determine if the timing of delivery and neonatal outcomes were significantly different between cohorts of mice, followed by *post-hoc* Bonferroni test for multiple comparisons. The level of significance was set at p≤0.05.

## 3. Results

### 3.1. Uterine-selectivity of mundulone and MA

As previously stated, off-target side effects of current off-label tocolytics on maternal vascular smooth muscle and the fetal DA have been major limitations of their use in the management of PL. In a prior study, we discovered that the transcriptomic profile of primary human aorta VSMCs was more similar to that of human primary myometrial cells, compared to human DA SMCs, thus serving as useful cells in assays to determine compound uterine-selectivity [41]. To this end, we used our previously established Ca^2+^-mobilization assay to examine whether mundulone and MA were uterine-selective based on either their lack of activity in VSMCs or greater effect on Ca^2+^-mobilization in myometrial SMCs compared to VSMCs. As shown in Fig.1 mundulone inhibited Ca^2+^-mobilization from myometrial cells and aorta VSMCs, while MA only inhibited Ca^2+^- mobilization in myometrial SMCs (Fig.1 A, C). Specifically, mundulone exhibited a significant (p<0.0001) 2- fold difference in potency, while MA displayed a significant (p=0.03) 70% difference in efficacy between myometrial cells compared to aorta cells (Fig.1B,D and Table 1). Furthermore, there was a significant difference between the efficacy (E_max_ = 80.5% vs. 44.5%, p=0.0005) and potency (IC_50_ = 27 μM vs. 14 μM, p=0.007) of mundulone and MA to inhibit Ca^2+^-mobilization in myometrial cells.

**Figure 1:**
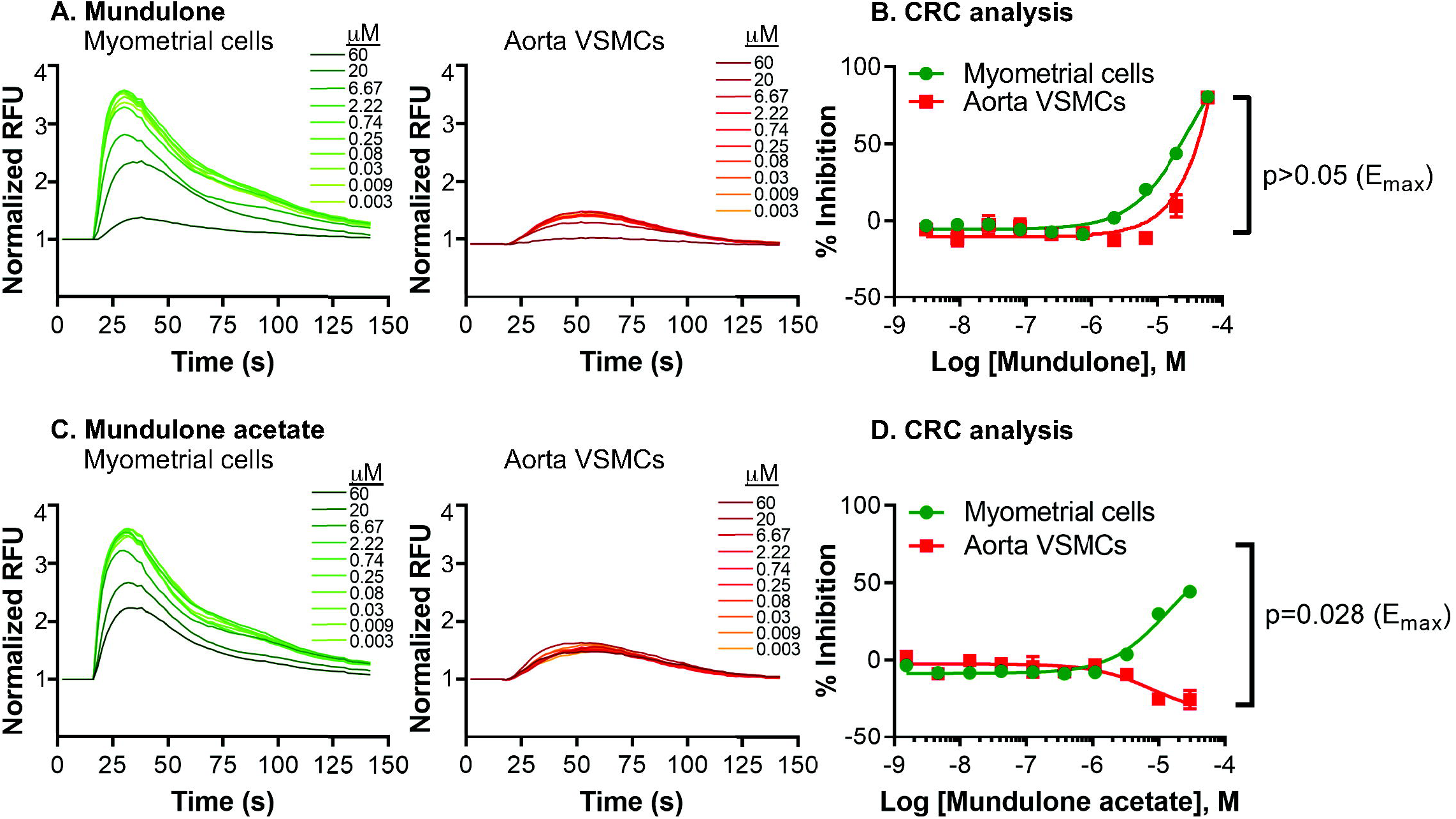
*In vitro* uterine-selectivity of mundulone and mundulone acetate (MA) to inhibit intracellular Ca^2+^- release. Real-time recording of concentration-dependent inhibition of intracellular Ca^2+^-release by mundulone (A) or MA (C) from myometrial cells and aorta VSMCs in 384-well format. Concentration-response curves of mundulone (B) or MA (D) showing selectivity towards myometrial cells compared to the aorta VSMCs due to either a significant shift in potency (IC_50_) or efficacy (E_max_), respectively. Non-linear regression was used to fit the data (mean ± SEM) and to calculate the IC_50_ and E_max_., which are provided in Table 1, along with p-values. A 2-way ANOVA with a post-hoc Fisher’s LSD test was used to compare the E_max_ values (shown).

**Table 1:**
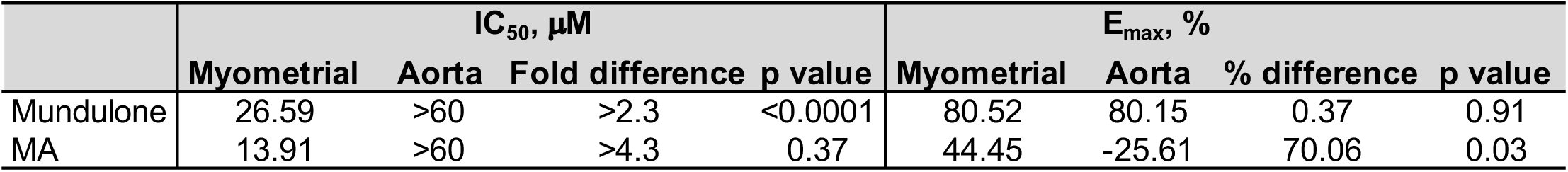
Uterine-selectivity of mundulone and mundulone acetate (MA)

### 3.2. *Ex vivo* tocolytic ability and lack of off-target effect on fetal DA

Established isometric contractility and vessel myography assays were used to examine the *ex vivo* uterine-selective effect of mundulone and MA on contractile activity of term pregnant myometrial tissue and the vasoreactivity of the term fetal DA, respectively [3, 55, 58–65].

As shown in Fig.2 A-B. mundulone, but not MA, displayed concentration-dependent inhibition of contractions of mouse uterine tissue (Fig.2B). The tocolytic efficacy and potency for mundulone (E_max_ = 70%, IC_50_=10 μM, p=0.02), but not MA (E_max_ = 27%, IC_50_>0.1 mM, p=0.25), was significantly different than the vehicle control. Mundulone exhibited less tocolytic efficacy (E_max_ = 57%) but similar potency (10 μM) to human compared to mouse myometrial tissue (Fig.2 B-D). Neither mundulone or MA affected the vessel diameter of the DA lumen beyond that of the vehicle control (∼10% difference from baseline; Fig.2 E-F, p>0.05).

**Figure 2:**
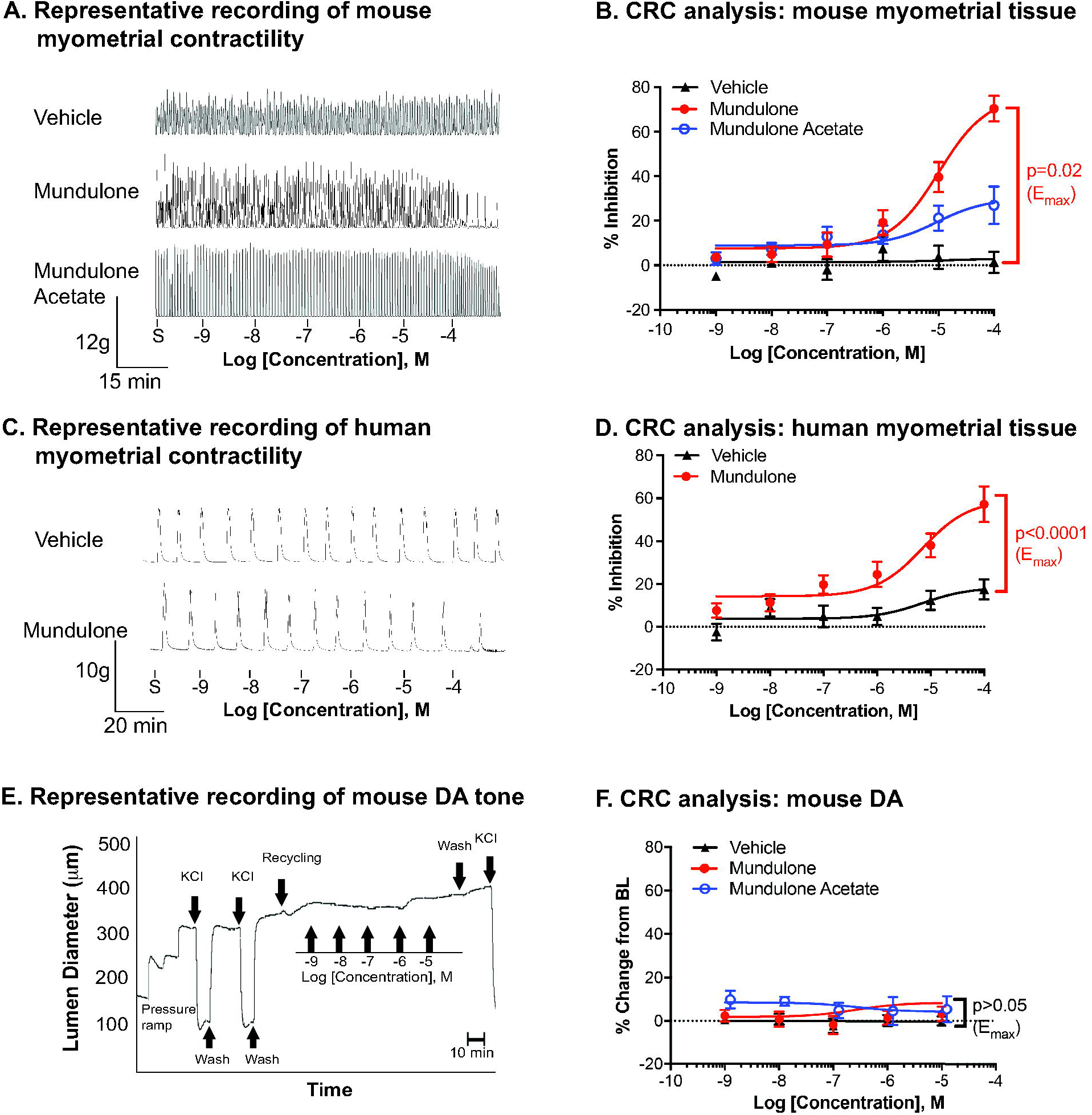
*Ex vivo* tocolytic ability and uterine-selectivity. Representative recording of isometric spontaneous contractility (measured in grams of tension) of mouse or human myometrial tissue (A and C, respectively) prior to the addition of increasing concentrations (10pm - 100μM) of vehicle control, mundulone or mundulone acetate (MA). Recordings were analyzed for contractile AUC (B and D). Non-linear regression was used to fit the data (mean ± SEM) and to calculate IC_50_ and E_max_. C. Representative tracing of fetal mouse ductus arteriosus (DA) tone (measured by lumen diameter) prior to and after the addition of increasing concentrations (1nm - 10μM) of vehicle control, mundulone or MA. D. Recordings were analyzed for % change from baseline lumen diameter. A 2-way ANOVA with a post-hoc Tukey analysis was used to compare the E_max_ values (shown).

### 3.3. *In vitro* synergism with current tocolytics

To improve the efficacy and/or potency of mundulone and MA, we explored the possibility of synergism when combined with current tocolytics. It is important to note that while these drugs function through different molecular targets, the exact mechanism(s) of action for mundulone and MA are currently unknown. We tested their synergistic potential in our *in vitro* high-throughput Ca^2+^-mobilization assay, which was adapted for compound combination-testing using 8x8 dose matrices (Fig.3A). We found that mundulone displayed synergism with atosiban and nifedipine (Fig.3B-C) at several concentration ratios, while MA exhibited synergistic efficacy with only nifedipine (Fig.3D). No synergy was observed in mundulone + indomethacin, MA + indomethacin and MA + atosiban combinations (Suppl Fig.1).

**Figure 3:**
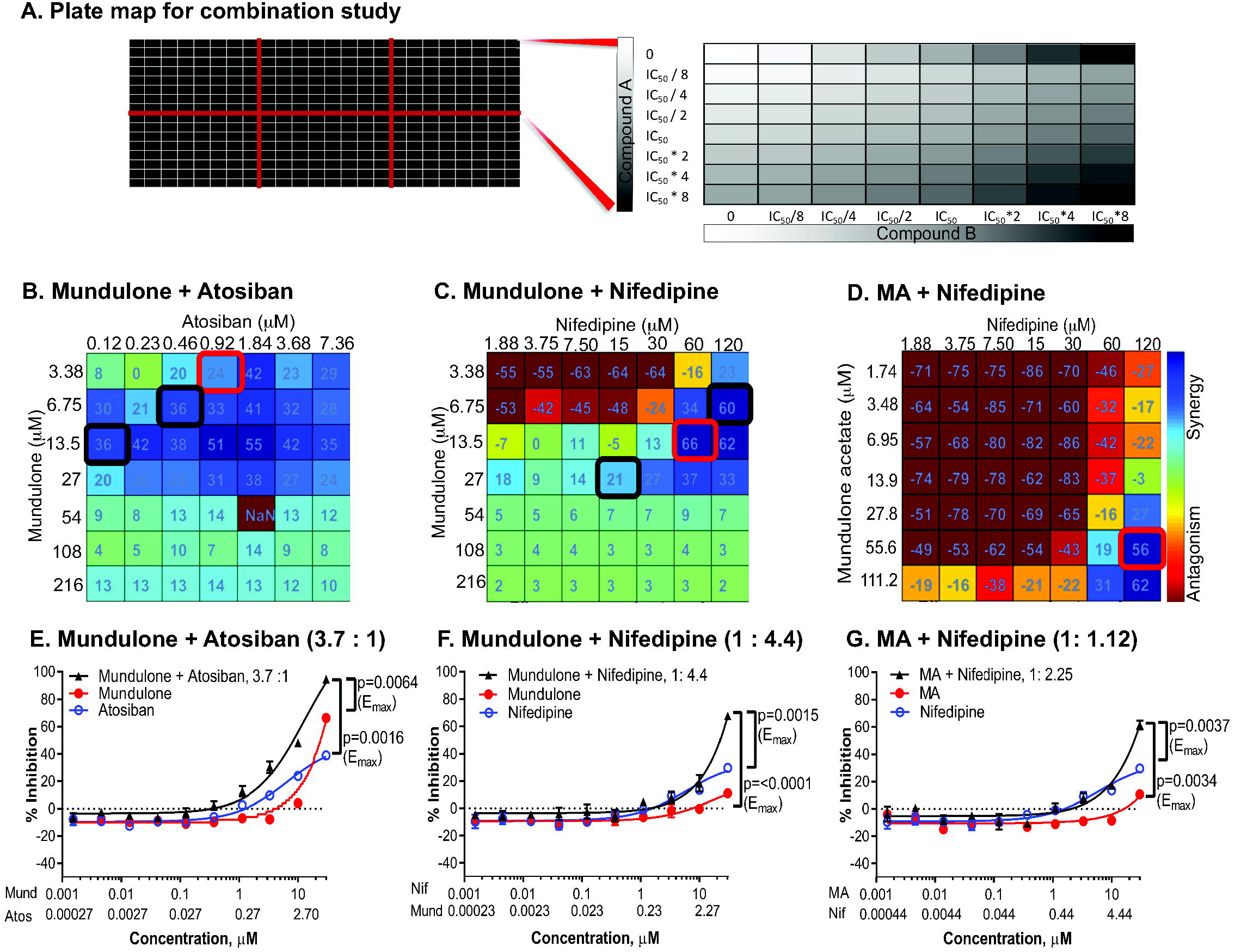
Identification of mundulone and mundulone acetate (MA) synergistic combinations with clinical tocolytics. A. Plate map used for the high-throughput combination Ca^2+^-assay in which one 384-well plate allows testing of six different combinations of two-compounds. Controls (no compounds, white squares) and individual compounds “A” (columns 1, 9 and 17) and “B” on (rows H and I, respectively) were included on each plate. The direction of the gradient shows the increasing concentrations of compound “A” and “B”. The %response data for each compound concentration was averaged and then analyzed with Combenefit software to provide synergy scores using the Bliss-independence model. Heat maps of synergy scores are shown (B-D). Red or black boxes indicate the three fixed ratios chosen for CRC analysis to confirmation synergistic potency or efficacy. Concentration-response curves of combinations at fixed ratio indicated in red box is shown (E-G). A 2-way ANOVA with a post-hoc Tukey analysis was used to compare the E_max_ values (shown).

For confirmation of synergistic efficacy and/or potency, we performed concentration-response analysis of FR of selected combinations (Fig.3E-G). The combination of mundulone + atosiban resulted in both increased efficacy and potency, while only increased efficacy was confirmed for mundulone + nifedipine and MA + nifedipine synergistic combinations. The E_max_ and IC_50_ obtained from CRCs of individual compounds in comparison with FRs of compounds in synergistic combinations are listed in Table 2, as are the results of their statistical comparisons. The potency of mundulone improved by a similar degree (2-3 fold) when in combination with either atosiban (FR 3.7:1) or nifedipine (FR 1:4.4) compared to mundulone as a single-agent. However, the efficacy of mundulone was further improved when in combination with nifedipine than atosiban (6-fold versus 2-fold, respectively), compared to mundulone as a single-agent. Mundulone and MA exhibited similar degrees of enhancement in potency (2-3 fold) and efficacy (6-fold) when in combination with nifedipine.

**Table 2:** Synergistic efficacy and /or potency of fixed-ratio combinations of mundulone and MA with current tocolytics.

| Cpd A + Cpd B | Ca<sup>2+</sup> assay, IC<sub>50</sub> (μM) |  |  | p value |  | Ca<sup>2+</sup> assay, E<sub>max</sub> (% inhibition) |  |  | p value |  |
| --- | --- | --- | --- | --- | --- | --- | --- | --- | --- | --- |
|  | Cpd A | Cpd B | Combo* | Combo vs Cpd A | Combo vs Cpd B | Cpd A | Cpd B | Combo | Combo vs Cpd A | Combo vs Cpd B |
| Mund + Atos, 112.5 : 1 | >30 | 7.4 | **>30; >0.27** | >0.05 | >0.05 | 66.3 | 4.55 | **93.66** | 0.0082 | <0.0001 |
| Mund + Atos, 14.7 : 1 | >30 | 12.99 | **25.95; 1.77** | >0.05 | >0.05 | 66.3 | 22.59 | **94.62** | 0.0016 | <0.0001 |
| Mund + Atos, 3.7 : 1 | >30 | 6.44 | **13.25; 3.58** | <0.0001 | 0.064 | 66.3 | 39.08 | **94.72** | 0.0064 | 0.0016 |
| Mund + Nif, 1 : 4.4 | 18.22 | 6.02 | >6.82; >30 | >0.05 | 0.0004 | 11.08 | 29.74 | **67.63** | <0.0001 | 0.0015 |
| MA + Nif, 1 : 2.25 | >30 | 6.02 | >13.33; >30 | >0.05 | 0.0015 | 10.68 | 29.74 | **61.47** | 0.0034 | 0.0037 |

Bolded values represent synergistic combinations in potency (EC<sub>50</sub>) and/or efficacy (E<sub>max</sub>). \*Values represent the concentration of compounds "A" and "B" present in the combination at 50% inhibition of Ca<sup>2+</sup>-mobilization. Cpd=compound, Mund=mundulone, Atos=atosiban, Nif=nifedipine, MA=mundulone acetate, Combo=combination

### 3.4. Cytotoxic effects on myometrial cells and metabolic organ cells

Prediction of *in vivo* cellular toxicity through *in vitro* cell viability assays using a suitable cell type is a key component in early drug discovery. To detect the likelihood of cellular toxicity in the target tissue (myometrium) and important xenobiotic metabolic organs (liver and kidney), we assessed the toxic effect of individual compounds mundulone and MA, as well as their synergistic combinations with current tocolytics, through a well-established WST-1 cell viability assay.

Mundulone had a significantly greater (p≤0.0005) effect on the viability of hTERT-HM, HepG2, and RPTEC cells compared to MA (Suppl. Fig.2 and Suppl. Table 1). In relation to the therapeutic benefit (*i.e.* potency of mundulone and MA in the Ca^2+^-assay), MA demonstrated a favorable *in vitro* TI =8.8, while the TI of mundulone was only 0.8. Suppl Table 1 lists the IC_50_ and TI for mundulone and MA for each cell type examined.

The FRs of synergistic combinations tested above were assessed for their effect on cell viability (Suppl. Fig.3A- C). Mundulone + atosiban was the only synergistic combination demonstrating a favorable TI = 10, which was a great improvement from the TI=0.8 of mundulone as a single agent. Unfortunately, the TI of MA in combination with nifedipine was much lower compared to MA as a single agent (1.8 vs 8.8, respectively). Table 3 lists the IC_50_ and TI for all compound combinations examined, as well as the IC_50_ for their respective single-drug controls.

**Table 3:** Therapeutic index of mundulone and MA in fixed-ratio combinations with current tocolytics.

| Drug combo | hTERT-HM |  |  |  | HEPG2 |  |  |  | RPTEC |  |  |  | TI |
| --- | --- | --- | --- | --- | --- | --- | --- | --- | --- | --- | --- | --- | --- |
|  | Cpd A | Cpd B | Combo* | p value | Cpd A | Cpd B | Combo* | p value | Cpd A | Cpd B | Combo* | p value |  |
|  | IC<sub>50</sub> (μM) | IC<sub>50</sub> (μM) | IC<sub>50</sub> (μM) |  | IC<sub>50</sub> (μM) | IC<sub>50</sub> (μM) | IC<sub>50</sub> (μM) |  | IC<sub>50</sub> (μM) | IC<sub>50</sub> (μM) | IC<sub>50</sub> (μM) |  |  |
| Mund + Atos, 3.7 : 1 | 58.9 | >200 | 133.9; 36.19 | 0.0038 | 22.1 | >200 | 132.4; 35.78 | <0.0001 | 26.02 | >200 | 169.2; 45.73 | <0.0001 | 10 |
| Mund + Nif, 1 : 4.4 | >200 | 106 | 19.15; 84.24 | <0.0001 | 92.6 | 150.2 | 23.16; 101.9 | 0.7152 | 119.5 | 143.9 | 40.25; 177.1 | 0.1978 | 2.81 |
| MA + Nif, 1 : 2.25 | 246.6 | 106 | 34.45; 77.52 | <0.0001 | 125.4 | 150.2 | 24.44; 55 | 0.0056 | >200 | 143.9 | 100; 225 | 0.0001 | 1.83 |

\*Values represent the concentration of compounds "A" and "B" present in the combination at 50% inhibition of Ca<sup>2+</sup> mobilization or cell viability, respectively. Bolded values represent a favorable therapeutic index (TI = IC<sub>50</sub> Cell Viability Assay/IC<sub>50</sub> Ca<sup>2+</sup> Assay). p-values provided for drug combinations are for comparison between the combination and compound A. Cpd=compound, Mund=mundulone, Atos=atosiban, Nif=nifedipine, MA=mundulone acetate, Combo=combination

### 3.5. *Ex vivo* tocolytic ability of synergistic combinations

Next, we determined whether the synergistic combination, mundulone + atosiban with a favorable TI, exhibited synergistic tocolysis on term pregnant mouse myometrium in an *ex vivo* contractility assay. The concentration-dependent inhibition of tissue contractility by the FR 3.7:1 of mundulone + atosiban was compared to that of the individual compounds (Fig.4). The efficacy and potency of mundulone to inhibit mouse myometrial contractions significantly (p=0.005) improved in combination with atosiban compared to use as a single agent (E_max_ = 97% and 0.09 μM versus E_max_ = 70% and IC_50_ = 10 μM, respectively; Fig.4A-B). Given this observation, we tested the tocolytic efficacy and potency of mundulone as a single-agent and in combination with atosiban using term-pregnant human myometrial tissue (Fig.4C-D). Similar results were yielded showing significantly increased efficacy (88% vs 57%, p=0.03) and potency (0.12 μM vs 7 μM, p=0.001) of mundulone in combination with atosiban compared to its use as single agent, respectively. The combination was confirmed to be synergistic for both mouse and human tissue (synergy score>10, Supplemental Table 2).

**Figure 4:**
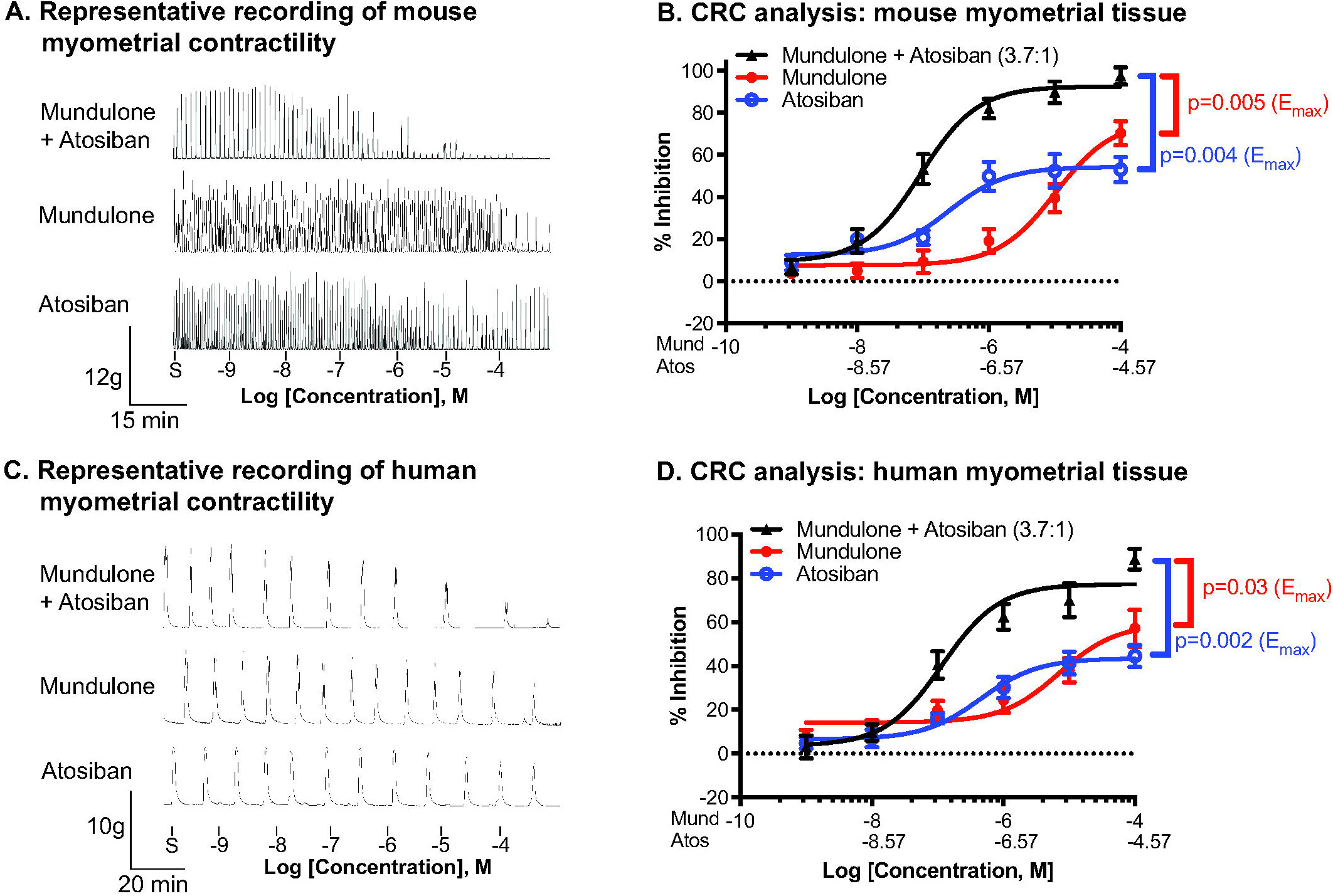
*Ex vivo* tocolytic effect of mundulone and atosiban combination. Representative recording of isometric spontaneous contractility (measured in grams of tension) of mouse or human myometrial tissue (A and C, respectively) prior to the addition of increasing concentrations (10pm - 100μM) of mundulone +atosiban at a fixed ratio (3.7:1), as well as their single-compound controls (mundulone or atosiban). Recordings were analyzed for contractile AUC (B and D). Non-linear regression was used to fit the data (mean ± SEM) and to calculate EC_50_ and E_max_. A 2-way ANOVA with a post-hoc Tukey analysis was used to compare the E_max_ values (shown).

### 3.6. *In vitro* ADME profiles

Mundulone and its acetate derivative were examined in a Tier 1 panel of ADME assays (Table 4). Both compounds were found to show low to moderate solubility and high permeability and were highly protein bound. Mundulone itself showed good stability in liver microsomes across all tested species, but the corresponding acetate was more labile, especially in rodent microsomes. This likely reflects instability of the acetate ester bond.

**Table 4:**
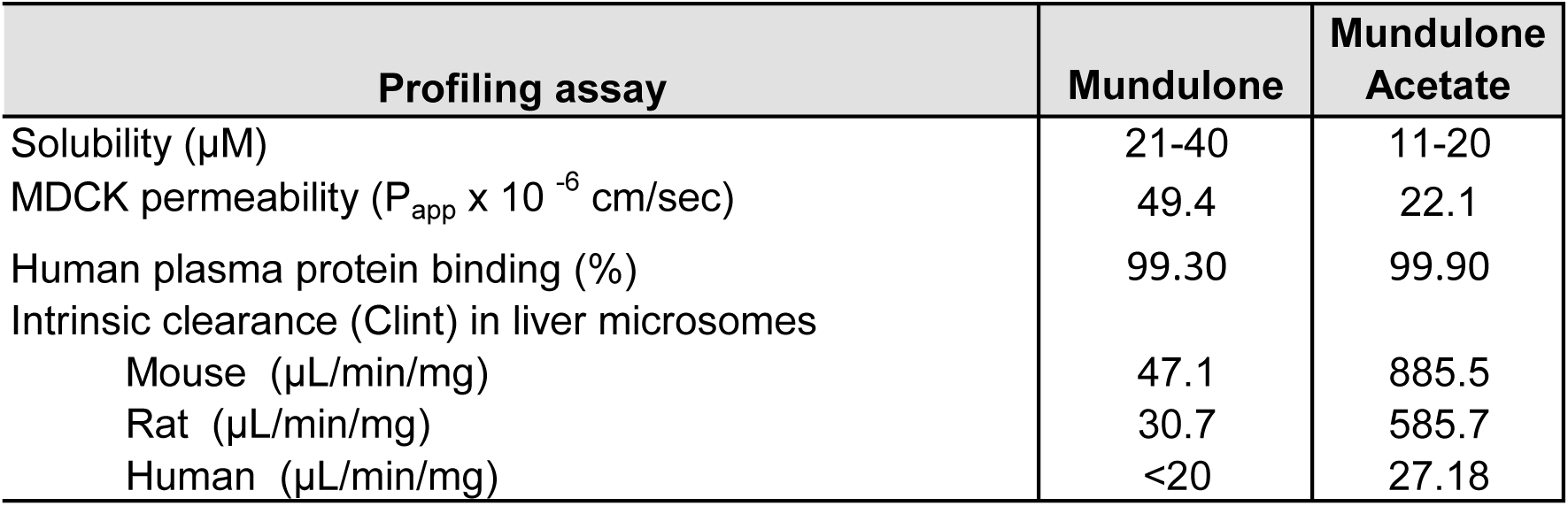
***In vitro* ADME profiles of Mundulone and Mundulone Acetate**

**Table 4:** Neonatal outcome measures.

| Treatment | Average |  | Surviving |  | Non-viable |  | Pup |  |
| --- | --- | --- | --- | --- | --- | --- | --- | --- |
|  | litter size | p value | pups born at term | p value | pups born at term | p value | weight at term delivery, g | p value |
| No treatments | 12.60 ± 3.11 |  | 12.5 ± 3.11 |  | 0.09 ± 0.29 |  | 1.64 ± 0.08 |  |
| MIF (no drug treatment) | 12.20 ± 1.64 | 1.00 | NA | NA | NA | NA | NA | NA |
| MIF + Mund + Atos, 6.5 + 1.76 mg/kg | 10.86 ± 2.55 | 0.46 | 9.60 ± 3.91 | 0.31 | 0.20 ± 0.45 | 0.87 | 1.55 ± 0.22 | 0.35 |
| MIF + Mund, 6.5 mg/kg | 12.00 ± 1.27 | 1.00 | NA | NA | NA | NA | NA | NA |
| MIF + Atos, 1.76 mg/kg | 12.00 ± 2.10 | 1.00 | 12.00 ± 0.00 | 1.00 | 0 | 0.99 | 1.65 ± 0.16 | 1.00 |
| MIF + Nif, 5 mg/kg | 11.90 ± 2.38 | 0.95 | 12.25 ± 3.86 | 1.00 | 0 | 0.87 | 1.66 ± 0.16 | 0.80 |

NA = Not applicable, no viable pups when delivered preterm. Maternal weights were only recorded until gestational day 18 or until delivery in cases of preterm birth. The data is represented as mean ± SD. Each group was compared to 'no treatment' group to determine if outcomes were significantly ( $p < 0.05$ ) different. MIF=mifepristone, Mund=mundulone, Atos=atosiban, Nif=nifedipine.

### 3.7. *In vivo* therapeutic efficacy to delay delivery and improve neonatal outcomes in a mouse model of PTB

Finally, the *in vivo* tocolytic efficacy and synergy of mundulone and atosiban in combination compared to the single agents were examined using a widely utilized mouse model of PTB for preclinical studies. We first determined the minimal effective dose of mifepristone to induce 100% PTB in mice, using the experimental design shown in Fig.5A. The timing of delivery of mifepristone did not become significantly (p≤0.0002) different than untreated control mice until 25 μg (Fig. 5B). The lowest doses required for 100% PTB rates were 30 μg of mifepristone (Fig. 5C).

**Figure 5:**
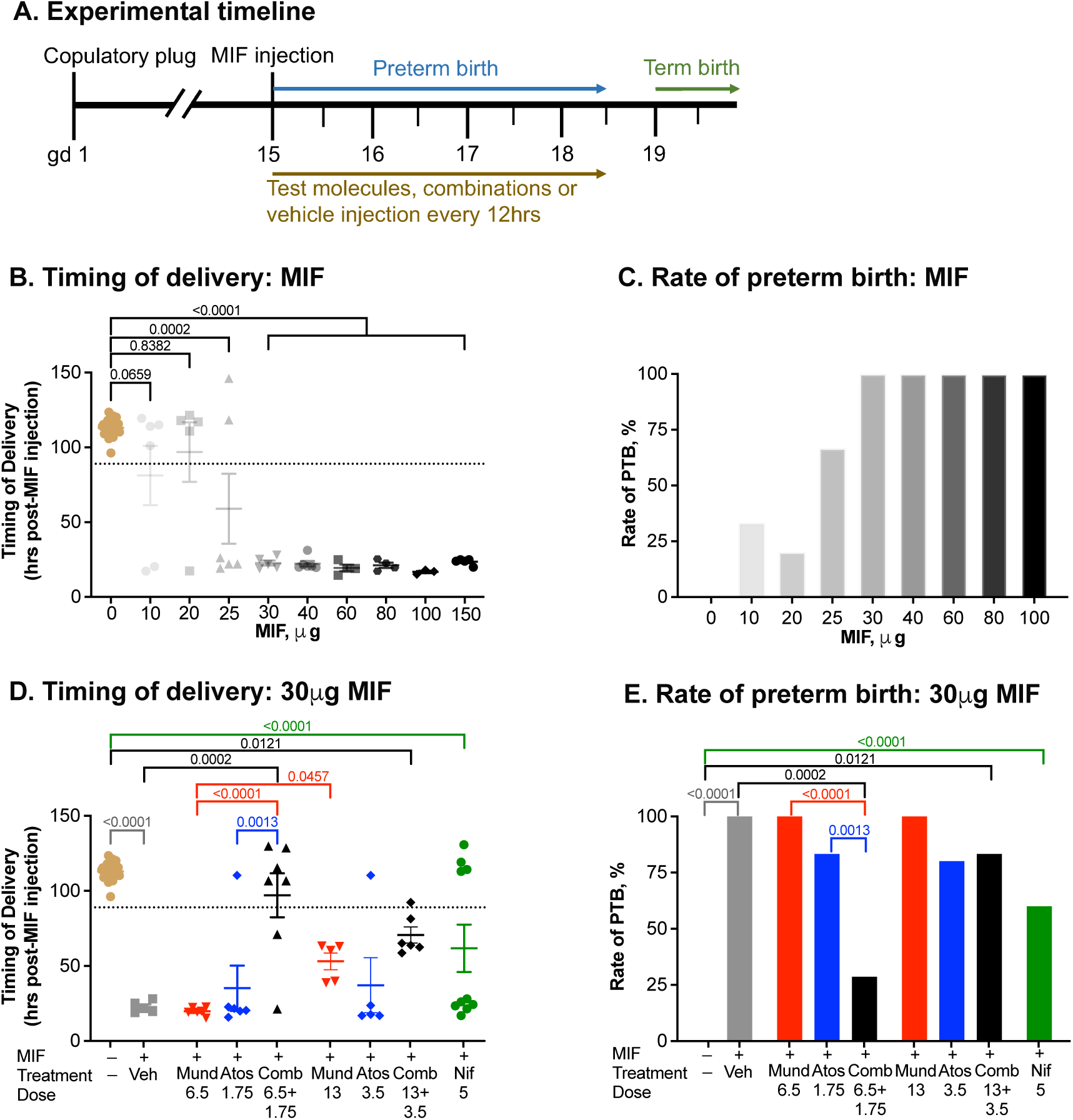
*In vivo* therapeutic efficacy of mundulone and atosiban combination to delay delivery in a preclinical mouse model of PTB. A. Experimental timeline for induction of preterm labor and birth using mifepristone (MIF) on day 15 of pregnancy, followed by administration of either vehicle, test or control drugs in mice. A. Determining the minimal effective dose of mifepristone to induce PTB in mice. Dose of mifepristone versus timing of delivery (B) or rate of PTB (C) in CD-1 pregnant mice. Timing of delivery (D) and PTB rates (E) for mice receiving either 30μg MIF (s.c.) prior to administration of either vehicle (Veh), mundulone + atosiban combination (Comb), or mundulone (Mund), or atosiban as single agents (Atos). Nifedipine (Nif) was used as a control tocolytic. Doses listed are mg/kg. Dotted line represents PTB, defined as 24hrs prior to average timing of term delivery of untreated mice.

Next, we assessed the efficacy of mundulone as a stand-alone agent and in combination with atosiban to delay mifepristone (30ug)-induced PTB in mice. Treatment with mundulone (6.5, 13, or 26 mg/kg twice daily) 5hrs after mifepristone administration (and PL induction) dose-dependently (p=0.0457) delayed the timing of delivery (Fig.5D). A 100% mortality was observed for mice treated with 26mg/kg mundulone twice daily until the evening of day 17. The mundulone+atosiban combination (FR 3.7:1, 6.5mg/kg + 1.75mg/kg) delayed the onset of delivery to term (>day19) for 71% of mice, with their timing of delivery occurring similarly (p>0.999) to non-mifepristone exposed mice (Fig.5E). None of the mifepristone-exposed mice treated with the combination displayed dystocia or post-term delivery. Only 17% and 0% of mice treated with atosiban (1.75mg/kg) or mundulone (6.5mg/kg) as single agents delivered at term, respectively. Interestingly, a higher dose of mundulone and atosiban (13mg/kg + 3.5mg/kg) at the same FR (3.7:1) resulted in only 17% of mice delivering at term, with 20% of atosiban (3.5mg/kg) and 0% of mundulone (13mg/kg) treated mice delivering at term. Nifedipine was used as positive control drug for these experiments, given its current clinical use as a tocolytic agent. Of the nifedipine-treated mice, 40% delivered at term, with their timing of delivery occurring significantly (p<0.0001) different from non-mifepristone exposed mice but statistically insignificant (p=0.999) than vehicle-treated mifepristone exposed mice. As shown in Table 4, the number of average surviving pups born at term was not significantly different across the groups. None of the pups born prematurely survived, thus their weights were not obtained. For the dams in each treatment group that were able to deliver at term, the average pup weights (as a ratio to the litter size) were not significantly different from that of the no treatment group.

Given the *in vivo* efficacy of the mundulone + atosiban (FR 3.7:1, 6.5mg/kg + 1.75mg/kg) to successfully delay delivery onset and manage PL in most mice exposed to 30 μg mifepristone, we next explored the therapeutic ability of this drug combination in the mifepristone model using the standard 150 μg dose, which we observed to be supramaximal for PTB-induction (Fig.5C). As shown in Supplemental Fig.4 A and B, mice exposed to 150 μg mifepristone that received treatment with the mundulone + atosiban combination delivered significantly (p=0.3) later than vehicle treated mice (52 hrs vs. 24hrs). While 28.6% of mice treated with the combination of mundulone and atosiban delivered at term, none of the nifedipine-treated 150 μg mifepristone mice delivered at term.

## 4. Comments

### 4.1. Principal findings

In the present study, we found that mundulone compared to MA displayed greater inhibitory efficacy on *in vitro* Ca^2+^-mobilization as well as *ex vivo* contractile activity. However, MA showed greater uterine-selectivity and a favorable *in vitro* TI, compared to mundulone, due to its lower cytotoxicity. A significant improvement of the TI was achieved through mundulone’s synergistic combination with atosiban, which yielded greater tocolytic efficacy and potency on term pregnant mouse and human myometrial tissue compared to single drugs. Furthermore, treatment with a combination of mundulone and atosiban in a 30 μg mifepristone-induced mouse model of PTB, significantly reduced the PTB rate compared to each single agent, allowing the dams to deliver at term-gestation without negative effects on pups.

Outside of its structural elucidation as an isoflavone [28, 66, 67], mundulone appears in only two publications to date, yielding little insight into its biological activity [68, 69]. In our study, mundulone and MA exhibited *in vitro* selectivity to inhibit Ca^2+^-mobilization from myometrial cells in comparison to aortic VSMCs, and *ex vivo* selectivity to inhibit myometrial contractility without observed effects on fetal DA vasoreactivity. While isoflavones are known to affect vascular [32, 70] and DA tissues[71] as well as uterine tissue [29], this study highlights that certain isoflavone structures can be uterine selective. Due to this, mundulone could benefit from medicinal chemistry efforts to study the structure-activity relationship and uterine-selectivity. Mundulone and its structurally-related derivative compound, dihydromunduletone, are reported to inhibit the G-protein coupled receptor GPR56’s activity [68]. Moreover, isoflavones are well known to be phytoestrogens with agonistic or antagonistic activity against estrogen receptors [72, 73]. Thus, it is worthwhile to probe the mechanism of both compounds, mundulone and MA, as it relates to modulation of intracellular Ca^2+^-regulated myometrial contractility through either a G-protein coupled receptor, estrogen receptor, or perhaps a different molecular target.

Past studies have examined whether combinations of tocolytics result in additive potency and/or efficacy [21–26]. The current study is the first to report a high-throughput combination screen using a dose matrix approach that includes a wide range of concentrations for the discovery of novel tocolytic synergy. We identified that the combinations of mundulone with two current tocolytics, atosiban and nifedipine, yielded *in vitro* synergistic effects on intracellular Ca^2+^-mobilization. Multiple pathways are involved upstream of intracellular-Ca^2+^ to regulate myometrial contractility. Moreover, the mechanism of action of mundulone to regulate uterine contractions remains unknown. Thus, the exact synergistic mechanism between mundulone-atosiban and mundulone-nifedipine remains to be determined, outside of our observation that the inhibition of Ca^2+^-channel as well as oxytocin receptor, but not the prostaglandin pathway, enhances the effect of mundulone. Our *in vitro* efficacy and combinational synergy of these novel molecules have been translated from the cell to the tissue level, as well as *in vivo*, thus, progressed through the discovery pipeline underscoring their translational potential.

When two drugs are combined, it is necessary to rule out unintended additive or synergistic toxicity early in the discovery process [74]. *In vitro* TI obtained by measuring a compound’s cytotoxicity is commonly used to quantify the extent of the safety at the desired efficacious conditions of a single- or combination of compounds. The toxicity of mundulone has been previously reported on HEK293 cells and zebrafish embryos [68, 75]; however, MA’s toxicity is not reported. While mundulone as a single-agent was found to be toxic at low concentrations (TI = 0.8) in our *in vitro* study, mundulone-treated mice did not show signs of impaired health or mortality until treated with 26mg/kg twice daily for two days. Nonetheless, mundulone could benefit from medicinal chemistry efforts to lessen its toxicity and improve its TI. In an alternative approach, we observed that the TI of mundulone dramatically improved to a favorable TI = 10 when combined with atosiban as a result of a 2-fold reduction in the concentration of mundulone, and the *ex vivo* tocolytic efficacy and potency significantly improved. To this end, at least one other study reported improved inhibition of uterine contractility when another flavonoid containing plant extracts from *Bryophyllum pinnatum* was combined with atosiban or nifedipine [36].

Currently, most test-compounds or drugs only significantly delay labor for a matter of hours using the mifepristone-induced mouse model of PTB [76–78]. However, there have been reports of a few investigational agents capable of managing PL for greater than 24hrs in the aforementioned mouse model, though not until term delivery [79–81]. To this end, it is possible that other test-compounds failed to manage PL until term delivery due to the commonly utilized dose (150μg) of mifepristone [82], which we found to be supramaximal to a 30μg dose that reliably induced 100% PTB rate in CD-1 mice. Indeed, nifedipine was able to delay delivery until term in 50% of PL mice induced with 30μg mifepristone, but 0% of mice receiving 150μg of the antiprogestin. The latter results are congruent with a prior study in which nifedipine resulted in delivery occurring within 25hrs post-mifepristone (150 μg) administration[76]. In our study, mundulone in combination with atosiban (FR 3.7:1, 6.5mg/kg + 1.75mg/kg) allowed long-term management of PL after induction with either 30 or 150 μg mifepristone, allowing 71% and 29% dams, respectively, to deliver at term (> day 19, 4-5 days post-mifepristone exposure) with viable pups without any visible maternal and fetal consequences. The efficacy of this drug combination outperformed barusiban and atosiban in animal models of preterm labor [83–85]. Additional preclinical studies are needed to determine the *in vivo* pharmacokinetics, biodistribution and placental transfer characteristics in mice and non-human primates.

There were limitations to the current study. First, the mouse model of PTB used in this study do not fully simulate the multifactorial causes of spontaneous PTB in humans, however it provides a “proof of principle” of the: (i) dose-dependent efficacy of mundulone as a single-agent to delay the timing of delivery and (ii) efficacy and synergy of the combination of mundulone with atosiban to manage PL until term delivery. The mifepristone mouse model of induced PL is a widely utilized model for preclinical testing of novel tocolytic agents for spontaneous PL in the absence of suspected or confirmed infection, and thus was selected for this study. However, it is important to examine the *in vivo* efficacy of mundulone as a stand-alone therapy or in combination with atosiban in additional established mouse models of PL using agents that either induce sterile inflammation (high-mobility group box-1 [86], interleukin-1 (alpha [87–89] and beta [89]), S100 calcium binding proteins-(A12 [90] and B [91]) or myometrial contractions (prostaglandin F2 alpha [92]) . Second, the long-term effects of *in utero* exposure to mundulone or MA was not examined, which would provide further insight into the neonatal safety of these compounds. Finally, while determining the target and mechanism of action of a newly identified molecule is important, it was beyond the scope of this study and will be evaluated in the future.

The present study demonstrates the tocolytic abilities of mundulone and MA through uterine-selective inhibition of intracellular-Ca^2+^. A novel *in vitro* high-throughput combination screen identified synergistic ratios of mundulone with atosiban with a favorable TI, whose tocolytic efficacy was validated in a separate *ex vivo* assay. Furthermore, the *in vivo* therapeutic efficacy and synergy of mundulone in combination with atosiban was ascertained in a mouse model of PTB. In summary, our findings highlight that mundulone or its analogs warrant future development as a stand-alone single- and/or combination-tocolytic therapy for management of PL.

## Acknowledgments

This project was supported by the *Eunice Kennedy Shriver* National Institute of Child Health and Human Development [Grants HD088830 and HD098213 (JLH), HD108420 (SS), and HD094946 (BCP)] and National Heart, Lung and Blood Institute Grants HL164327 and HL128386 (JR). We are thankful to the nurses and physicians in the Department of Obstetrics and Gynecology at Vanderbilt University Medical Center for collecting myometrial biopsies, and the women that kindly participated in this study. We thank the Vanderbilt Institute of Chemical Biology High Throughput Screening Facility for their technical assistance and use of their equipment. The Wave Front Biosciences Panoptic was purchased with funds from the NIH Office of The Director S10OD021734. We thank Dr. Jennifer Condon (Wayne State University) for kindly providing the hTERT-HM cells used in this study. All authors have read the journal’s authorship agreement and policy on disclosure of potential conflicts of interest.

---

Supplementary Figure 1. Combinations of mundulone and MA with clinical tocolytics that were not synergistic. A high-throughput Ca^2+^-assay using myometrial cells and mundulone or MA in combination with clinical tocolytics, indomethacin or atosiban, was performed. The %response data for each compound concentration was averaged and then analyzed with Combenefit software to provide synergy scores using the Bliss-independence model. Heat maps of synergy scores are shown for combinations of mundulone and MA with clinical tocolytics that did not display synergy: mundulone+indomethacin (A), MA+atosiban (B) and MA+indomethacin (C).

Supplementary Figure 2. Effect of mundulone and MA on cell viability. A WST-1 assay was used to examine the % cell viability of myometrial (hTERT-HM) cells, liver (HepG2) cells and kidney (RPTEC) cells after 72hr incubation with either mundulone, MA, DMSO (vehicle control) or napabucasin (positive control, known-toxic compound). Non-linear regression was used to fit the data (mean + SEM) and calculate IC_50_, which are provided in Suppl. Table 1, along with p-values and therapeutic indices.

Supplementary Figure 3. Cytotoxicity assessment of mundulone and mundulone acetate (MA) synergistic combinations with clinical tocolytics. A WST-1 assay was used to examine the % cell viability of myometrial (hTERT-HM) cells, liver (HepG2) cells and kidney (RPTEC) cells after 72hr incubation with synergistic combinations of mundulone + atosiban (A), mundulone + nifedipine (B) and MA + nifedipine (C) at fixed ratios indicated on the graphs, and their respective single-compound controls (Mund = Mundulone, Nif = Nifedipine, Atos = Atosiban, and MA). Non-linear regression was used to fit the data (mean ± SEM) and calculate IC_50_, which are provided in Table 3, along with p-values. A 2-way ANOVA with a post-hoc Tukey analysis was used to compare the E_max_ values (shown).

Supplemental Figure 4. I*n vivo* therapeutic efficacy of mundulone and atosiban combination to delay delivery in the commonly-utilized dose (150μg) of MIF-induced PTB model. Timing of delivery (A) and PTB rates (B) for mice receiving 150μg MIF (s.c.) prior to administration of either vehicle (Veh), mundulone + atosiban combination (Combo; 6.5 + 1.76mg/kg), or nifedipine (Nif; 5mg/kg). Dotted line represents PTB, defined as 24hrs prior to average timing of term delivery of untreated mice.

