## Supplemental Table 1 for "Arrest of mouse preterm labor until term delivery by combination therapy with atosiban and mundulone, a natural product with tocolytic efficacy"

**Supplemental Table 1: Therapeutic index of mundulone and MA as single-agents**

| | $Ca^{2+}$ assay | Cell viability assay | | | | | | |
|---|---|---|---|---|---|---|---|---|
| | Myometrial | hTERT-HM | | HEPG2 | | RPTEC | | TI |
| **Single-agent** | **$IC_{50}$ (μM)** | **$IC_{50}$ (μM)** | **p value** | **$IC_{50}$ (μM)** | **p value** | **$IC_{50}$ (μM)** | **p value** | |
| Mundulone | 26.59 | 58.9 | 0.0005 | 22.1 | <0.0001 | 26.02 | <0.0001 | 0.8 |
| MA | 13.91 | 122.7 | | 124.3 | | >200 | | **8.8** |

Bolded values represent a favorable therapeutic index (TI = $IC_{50}$ Cell Viability Assay/$IC_{50}$ $Ca^{2+}$ Assay).
