## Supplemental Table 2 for "Arrest of mouse preterm labor until term delivery by combination therapy with atosiban and mundulone, a natural product with tocolytic efficacy"

**Supplementary Table 2: Synergy scores for mundulone and atosiban combination on *ex vivo* myometrial contractility**

| Mundulone + Atosiban, 6.5 + 1.76 mg/kg | Synergy score | |
|---|---|---|
| | Bliss | HSA |
| Human | 14.85 | 14.58 |
| Mouse | 23.07 | 25.68 |

Synergy score >10 indicate synergism
