## Supplementary figures and images for "Arrest of mouse preterm labor until term delivery by combination therapy with atosiban and mundulone, a natural product with tocolytic efficacy"

### Supplemental Figure 1

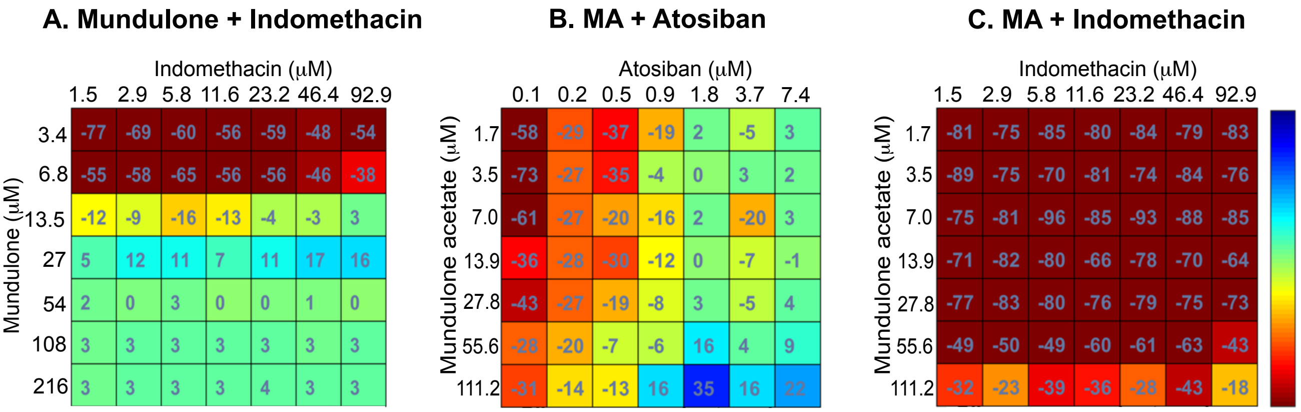

### Supplemental Figure 4

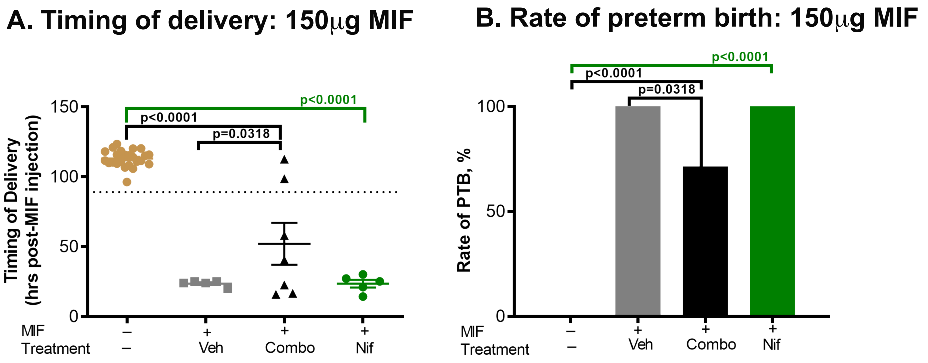
