## Supplemental Figure 2 for "Arrest of mouse preterm labor until term delivery by combination therapy with atosiban and mundulone, a natural product with tocolytic efficacy"

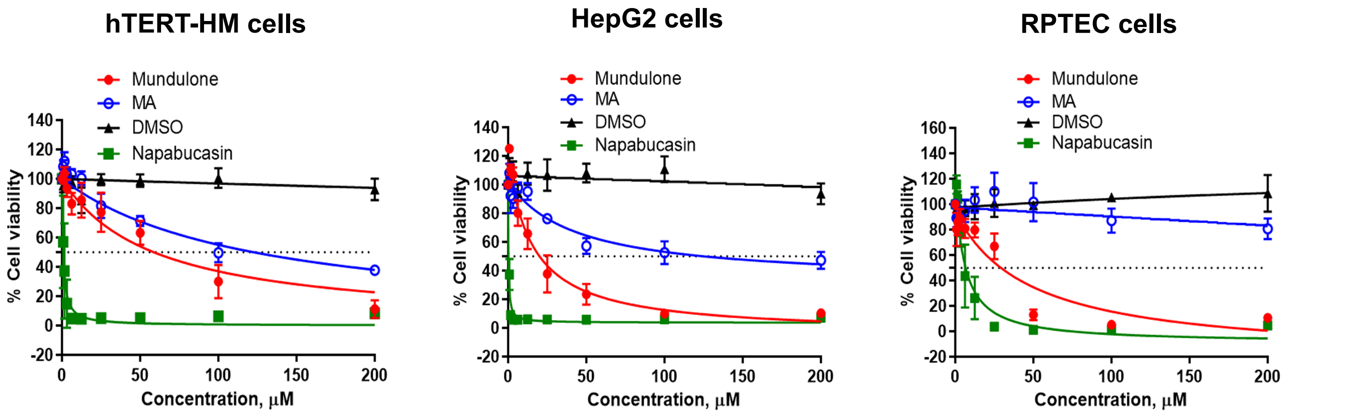


**Supplementary Figure 2.** Effect of mundulone and MA on cell viability. A WST-1 assay was used to examine the % cell viability of myometrial (hTERT-HM) cells, liver (HepG2) cells and kidney (RPTEC) cells after 72hr incubation with either mundulone, MA, DMSO (vehicle control) or napabucasin (positive control, known-toxic compound). Non-linear regression was used to fit the data (mean + SEM) and calculate IC50, which are provided in Suppl. Table 1, along with p-values and therapeutic indices.
