## Supplemental Figure 3 for "Arrest of mouse preterm labor until term delivery by combination therapy with atosiban and mundulone, a natural product with tocolytic efficacy"

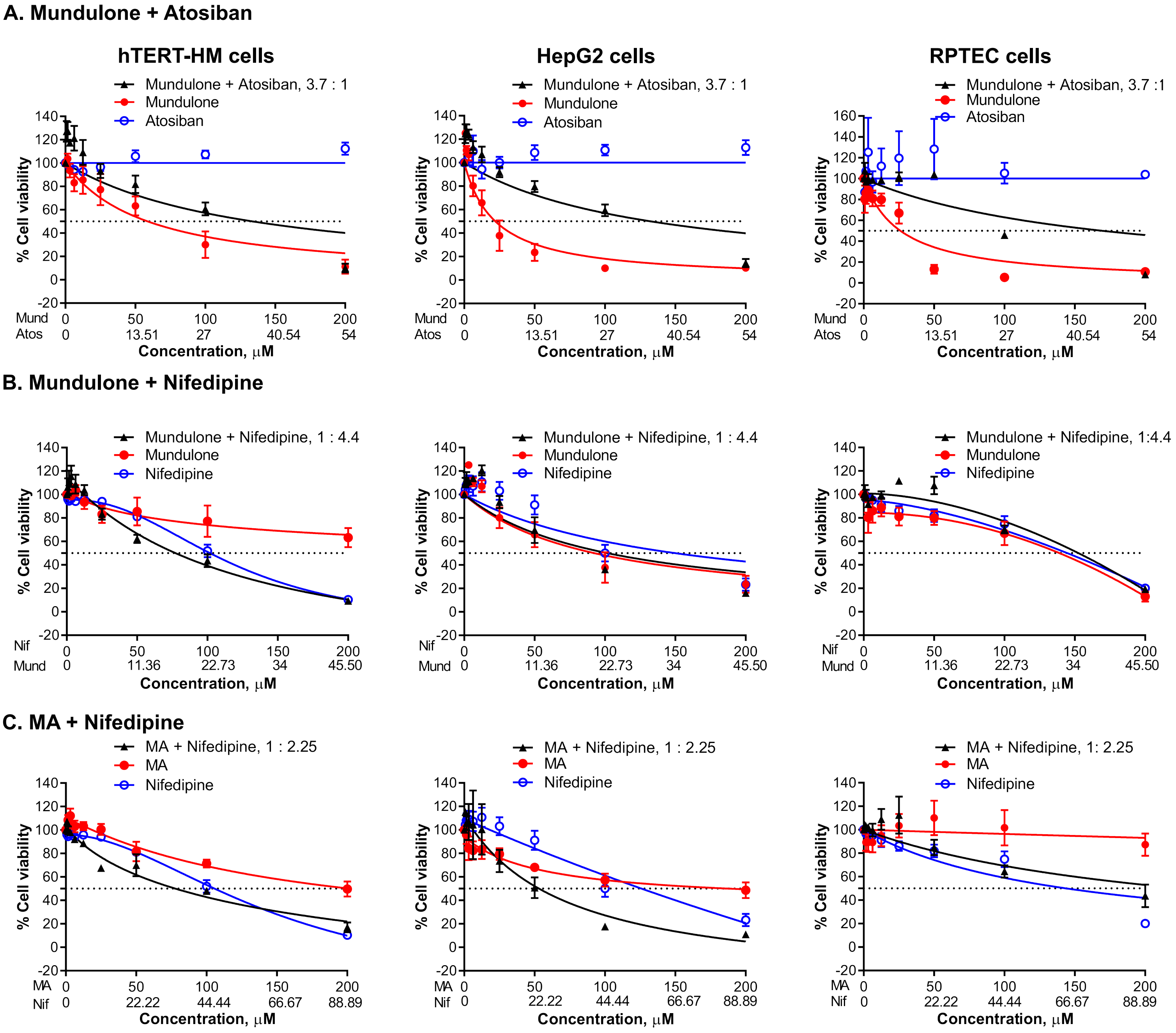


**Supplementary Figure 3.** Cytotoxicity assessment of mundulone and mundulone acetate (MA) synergistic combinations with clinical tocolytics. A WST-1 assay was used to examine the % cell viability of myometrial (hTERT-HM) cells, liver (HepG2) cells and kidney (RPTEC) cells after 72hr incubation with synergistic combinations of mundulone + atosiban (A), mundulone + nifedipine (B) and MA + nifedipine (C) at fixed ratios indicated on the graphs, and their respective single-compound controls (Mund = Mundulone, Nif = Nifedipine, Atos = Atosiban, and MA). Non-linear regression was used to fit the data (mean ± SEM) and calculate IC50, which are provided in Table 3, along with p-values. A 2-way ANOVA with a post-hoc Tukey analysis was used to compare the Emax values (shown).
